# West Nile virus is transmitted within mosquito populations through infectious mosquito excreta

**DOI:** 10.1101/2024.01.29.577888

**Authors:** Rodolphe Hamel, Quentin Narpon, Idalba Serrato-Pomar, Camille Gauliard, Arnaud Berthomieu, Sineewanlaya Wichit, Dorothée Missé, Mircea T. Sofonea, Julien Pompon

**Affiliations:** MIVEGEC, Univ. Montpellier, IRD, CNRS, Montpellier, France; Department of Clinical Microbiology and Applied Technology, Faculty of Medical Technology, Mahidol University, Bangkok, Thailand; Viral Vector Joint Unit and Join Laboratory, Mahidol University, Thailand; IHU Méditerranée Infection, Marseille, France; PCCEI, Univ. Montpellier, Inserm, EFS, Montpellier, France; CHU Nîmes, Nîmes, France

## Abstract

Understanding transmission routes of arboviruses is key to control their epidemiology and global health burden. Using West Nile virus and *Culex* mosquitoes, we tested whether arboviruses are transmitted through mosquito excreta. First, we determined the presence of infectious virions in excreta and quantified a high concentration of infectious units per excreta. Second, we showed that virion excretion starts early after oral infection and remains constant for a long period, regardless of mosquito infection level. These results highlight the infectiousness of excreta from infected mosquitoes. Third, we found that both larvae and pupae were susceptible to infection, although pupae were highly permissive. Forth, we established the proof-of-concept that immature mosquitoes can be infected by infectious excreta, demonstrating a new excreta-mediated mode of transmission. Finally, by mathematically modelling excreta-mediated transmission in the field, we demonstrated its potential impact on arbovirus epidemiology. Our study uncovers a new route of transmission for arboviruses, unveiling mechanisms of viral maintenance in mosquito reservoirs and of vector species shift that contribute to zoonotic emergence.

## Introduction

Originally isolated in the West Nile province of Uganda in 1937 [1], West Nile Virus (WNV) is currently the most widely distributed mosquito-borne diseases [2]. Circulation of WNV has been reported on all continents, except Antarctica [3–6]. Although WNV infection in humans remains asymptomatic in most cases, approximately 25% of infected patients develop non-lethal flu-like symptoms and 1% show neurological manifestations such as encephalitis, meningitis, or acute flaccid paralysis, potentially causing death and long-term sequelae [7]. Furthermore, an epidemiologic shift in the 90’s resulted in increased severity of outbreaks with more frequent neurological symptoms [8]. Initially observed around the Mediterranean basin, the more virulent lineage 1 was introduced in the USA in 1999 and rapidly spread throughout the country and the Americas. Since 2000, WNV infected an estimated 7 million people and caused more than 2,700 deaths in the USA [9,10], while the disease causes yearly deaths in the EU where more than 100 people died in 2022 and 2023 [11]. Despite the alarming situation, there is neither therapeutics nor licensed vaccines for humans [3,6].

WNV transmission occurs through multiple routes. Primarily, WNV is transmitted between vertebrate hosts through mosquito vectors, mostly from the *Culex* genera; a mode that is referred to as “horizontal” transmission [2]. Successful horizontal transmission occurs when a susceptible mosquito bites an infected host. The virus then multiplies within the vector until infecting the salivary glands, from which it is expectorated in the skin of other susceptible hosts during subsequent blood feeding, resulting in transmission [12]. WNV circulates in an enzootic cycle between birds, where *Cx. quinquefasciatus*, *Cx. pipiens* and *Cx. tarsalis* are the main vectors. Occasionally, opportunistic feeding of some *Culex* species result in transmission from birds to humans or other mammals [13,14]. However, mammals are dead-end hosts as most of them do not develop a sufficiently high viremia to infect mosquitoes. Additionally, WNV can be directly transmitted between vertebrate hosts by contact with or consumption of infectious material, such as infected birds, mosquitoes, cloacal fluids, blood transfusion, organ transplantation or even breast milk [15–17]. Finally, WNV as for other flaviviruses can be maintained within mosquito populations by direct transmission from an infected female mosquito to its offspring; a mode referred to as “vertical” transmission [18–20]. However, low vertical transmission rates reported in laboratories imply a moderate epidemiological role [21], even though vertical transmission efficiency improves with extrinsic incubation duration [18,22].

Several lines of evidence indicate that WNV is maintained within mosquito populations without cycling through vertebrate hosts. WNV was detected in *Culex* males [23], in larvae [24,25] and pupae [26], all of which became infected by exposition to another inoculum source than blood. Circulation of the virus between mosquitoes then enables persistence of the virus when conditions are unfavorable for horizontal transmission, and facilitate resurgence of transmission to vertebrate, including humans, when conditions favor mosquito biting to susceptible hosts [8,27]. Understanding the modes of transmission that maintain viruses within mosquito populations is important to promote novel interventions and improve epidemiological forecast to adjust interventions.

Here, we test whether WNV can be maintained in mosquito populations through excreta-mediated transmission. Our hypothesis is based on the observation that excreta from infected mosquitoes contain detectable amount of arboviral RNA and for this reason are screened as an innovative surveillance strategy [28,29]. Furthermore, a previous study observed that excreta from *Cx. annulirostris* mosquitoes carry infectious WNV virions but concluded that the amount was too low to infect other mosquitoes [29]. In our study, we used WNV as a flavivirus model and showed that infected *Cx. quinquefasciatus* excrete infectious virions. We then evaluated the possibility of an excreta-mediated transmission to immature mosquitoes by: (i) quantifying the inoculum per excreta; (ii) assessing how extrinsic incubation period and mosquito infection intensity influence excreta infectivity; (iii) determining the susceptibility of immature mosquitoes to viral infection; and (iv) demonstrating that infectious excreta can infect immature mosquitoes. Eventually, we combined our multifactorial dataset into a mathematical model to assess the potential for excreta-mediated WNV transmission in breeding sites. Our study uncovers a new mode of transmission for WNV and probably all arboviruses through infectious excreta, improving our understanding of arbovirus epidemiology.

## Materials and Methods

### Cells, viruses, and mosquitoes

C6/36 cells (ATCC CRL-1660) derived from *Aedes albopictus* and Vero cells (ATCC-CCL-81) derived from green monkey (*Chlorocebus sabaeus*) kidney were grown in Dulbecco’s modified Eagle’s medium (DMEM) (Invitrogen, France) supplemented with 10% fetal bovine serum (FBS) (Eurobio, France), 1% penicillin-streptomycin (Gibco, France). Insect cells medium was also supplemented with 1 % non-essential amino acid (Gibco, France). Mosquito cells were grown at 28°C and mammalian cells at 37°C, while both cells were grown with 5% CO_2_.

A WNV infectious clone derived from IS98, a highly virulent strain isolated from a white stork, *Ciconia ciconia* (IC-WNV-IS98; Genbank accession number: KR107956.1), was received from Dr. Philippe Desprès [30] and propagated in C6/36 cells before storage at -70°C.

*Culex quinquefasciatus* strain SLAB originating from California were bred in the Montpellier Vectopole Sud facility. Larvae were maintained in plastic trays (Gilac^®^, France) with distilled water and fed a mixture of pelleted rabbit food (Hamiform™, France) and fish TetraMin flake (Tetra^®^, France). L1 larvae were also initially given yeast solution. Pupae were transferred to a new tank and placed in a net cage (29 x 18 x 22 cm) (Custom manufacturing) with water and sugar solution (10%) for emerging adults. Mosquitoes were maintained at 26-28°C, 70-80% humidity with a 12h:12h photoperiod.

### Oral infection

Adult mosquitoes aged 3 to 5 day-old were sedated at +4°C, sorted at a density of 50 females and 5 males per box and starved for 24h. Mosquitoes were then transferred to the BSL3 insectary to acclimatize at 28°C with 80% humidity for 3 hours. Hemotek^®^ membrane feeding system (Hemotek Ltd, United Kingdom) was used for oral infection using chicken’s skin and an infection mixture consisting of 1,500 µl PBS-washed-rabbit blood (IRD animal facility, accreditation number H3417221), 150 µl FBS, 150 µl of 5 mM ATP (Sigma-Aldrich, France), 700 µl Roswell Parc Memorial Institute medium (RPMI) (Gibco, France) and WNV stock to obtain either 10^5^ or 10^7^ pfu/ml of blood. Mosquitoes were allowed to feed on the blood mixture maintained at 37°C for 1h15. Fully engorged mosquitoes were then sorted in an appropriate container with *ad libitum* access to water and sugar solution (10%).

### Collection of excreta

To avoid detecting viruses secreted during feeding on the WNV-blood meal, mosquitoes were transferred into new containers 2-3 days post exposure (DPE), when the blood was digested. Different types of containers were used for collecting pooled or single excreta.

For pooled excreta collections, at 6 DPE, female mosquitoes were grouped in 250 mL jars (Nalgene, France) at a density of 25 mosquitoes/jar. Mosquitoes were offered sugar solutions (10%) containing a blue food colorant (Vahiné, France). Excreta were then collected over intervals of 1h-1h30 in 500 µl of DMEM containing 1% antibiotic-antimycotic (Gibco, France). Before adding media, the number of excreta was visually counted as blue dots.

For single excreta collections, female mosquitoes were maintained in round-bottomed 14 mL polypropylene Falcon tubes (Fisher Scientific, France), crowned with a cap manufactured by a 3D printer to allow mosquito feeding on a sugar solution (10%) and safe mosquito transfer from one tube to another to collect excreta without sedating mosquitoes (Sup. Fig. 1). Excreta were collected in 500 µl of DMEM containing 1% antibiotic-antimycotic (Gibco, France) on the 4^th^, 6^th^, 8^th^, 10^th^ and 12^th^ DPE. On the twelfth day, mosquitoes were collected and analyzed. During excreta collection, mosquitoes were maintained in a climatic chamber at 28°C, 80% humidity and a 12h:12h photoperiod.

### Infection of cells with excreta

Media containing pooled excreta was filtered through 0.22 µm filter (Milex-GV^®^, Fisher Scientific, France) and 150 µl of the filtrate were combined with 350 µl of DMEM to inoculate T25 flasks containing 8.5 x 10^5^ Vero cells for 1h15 at 37°C. After washing, cells were incubated for 6 days at 37°C with 5% CO_2_. Supernatant was collected, filtered (filter exclusion size 0.45µm, Fisher Scientific, France) and analyzed by RT-PCR and plaque assay.

### RNA extraction

Single adult mosquitoes were homogenized in a 1.5 ml Eppendorf tube with plastic pestle in 500 µl of TRI Reagent (Euromedex, France) before RNA extraction according to manufacturer’s instructions. Single larval and pupal mosquitoes were similarly homogenized in 500 µl of TRI Reagent before RNA extraction according to manufacturer’s instructions. RNA from 150 µl of excreta solution was extracted by adding 600 µl of RAV1 lysis buffer and using NucleoSpin virus RNA kit (Macherey-Nagel, France).

### WNV gRNA detection by RT-PCR and quantification by RT-qPCR

RT-PCR was performed using AccessQuick RT-PCR System (Promega, France) in total reaction volume of 25 µl with 5 µl of RNA extracts and 400 nM of forward primer (5’-ATTCGGGAGGAGACGTGGTA-3’) and reverse primer (5’-CAGCCGCCAACATCAACAAA-3’) to amplify a 129 base pairs (bp) in the WNV envelope region. Reactions were conducted at 42°C for 45 min, 95°C for 2 min followed by 45 cycles of 20s at 95°C, 20s at 58°C and 20s at 72°C and a 2 min-final step at 72°C. PCR products were visualized on 2% agarose gel.

One-step RT-qPCR was conducted using GoTaq 1-Step RT-qPCR System kit (Promega, France) in total reaction volume of 20 µl containing 2 µl of RNA extracts and 300 nM of the same forward and reverse primers as above. Amplification was conducted on AriaMax Real-Time PCR system (Agilent, France) and consisted of an initial RT step at 42°C for 20 min, 95°C for 10 min, followed by 45 cycles of 10s at 95°C, 15s at 60°C and 20s at 72°C, and a final melting curve analysis. Viral RNA was absolutely quantified by establishing a standard equation using serial dilutions of known amounts of the *in vitro* transcribed qPCR RNA target. The amplicon target was amplified from WNV cDNA using the qPCR primers with the forward primer flanked by T7 sequence (5’-TAATACGACTCACTATAGGGATTCGGGAGGAGACGTGGTA-3’) and transcribed using T7 RiboMAX Express Large Scale RNA Production System kit (Promega, France). RNA was purified by ethanol precipitation, quantified by NanoDrop spectrophotometer (FisherScientific, France) and converted to concentration of molecular copies by using the following formula: number of Viral RNA copy / µl = [(g/µl of RNA)/(transcript length in bp x 340)] x 6.02 x 10^23^.

### WNV titration

Triplicates of 1.8 x 10^5^ Vero cells were infected with 10-fold serial dilutions of 250 µl of excreta solution or cell supernatant at 37°C for 1h15. After washing, cells were overlaid with DMEM containing 2% carboxymethylcellulose (CMC, Sigma-Aldrich, France), 2% FBS and 1% of antibiotic-antimytotic (Gibco, France). Cells were incubated at 37°C with 5% CO_2_ for 7 days. The overlay medium was then aspirated, and cells were incubated 30 min at room temperature with 3.7% formaldehyde diluted in PBS, washed twice with PBS, and incubated with crystal violet solution (3.7% formaldehyde and 0.1% crystal violet in 20% ethanol) for 1h. After two washes, plaques were counted and used to calculate PFU/ml with the following formula:

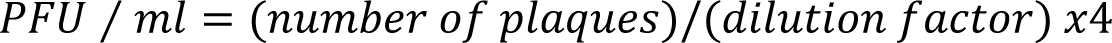

### WNV stability

5 x 10^4^ PFU/ml of WNV was incubated in water supplemented with larval food at 28°C. 200 µL of liquid were collected after 0 min, 30 min, 1h, 2h and used for viral titration.

### Infection of mosquito aquatic stages

Fifteen L1 *Cx. quinquefasciatus* larvae were incubated for 1h in one Petri dish (Nunclon™, FisherScientific, France) containing 2 ml of food-supplemented water and different concentrations of WNV stock. Larvae were then transferred to plastic tubes (Nalgen, France) capped with cotton and containing 3 ml of distilled water with larval nutrient solution and incubated at 28°C, 80% humidity. On day 5 post exposition, L4 larvae were collected, rinsed twice in distilled water and collected for RNA extraction. Twenty-five pupae were similarly incubated with WNV and, after exposure, were transferred inside a rearing cage and kept at 28°C, 80% humidity with sugar solution (10%). RNA extraction was performed on adult mosquitoes collected three days after emergence.

Seven and eight pupae were separately incubated with 300 µl of pooled excreta solution in one well of 48-well flat-bottom plate (Falcon™, Fisher Scientific, France). Pupae were placed in a climatic chamber with rearing conditions. Adults were collected in 500 µl of TRI Reagent for RNA extraction three days after emergence.

### Mathematical modeling of stercoraceous transmission

A mathematical model governed by an autonomous non-linear dynamical system governed by five ordinary differential equations (ODE) (see below) was analyzed through the next-generation theorem [31] to derive a closed-formed expression of the basic reproduction number for transmission through mosquito excreta, *R*^d^ (see below).

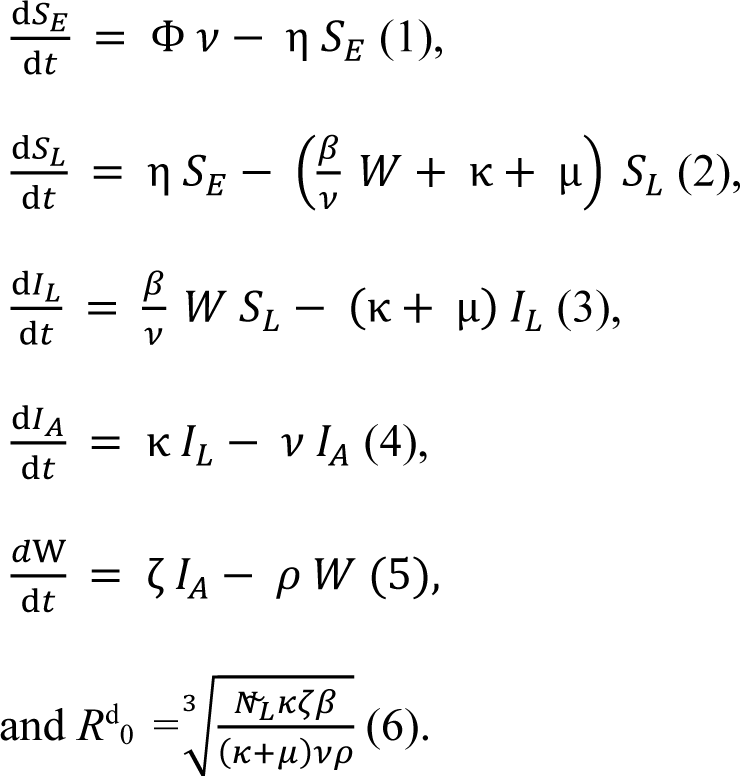

Where S_E_ stands for surviving eggs; S_L_ immature mosquito susceptibility; I_L_ infected immature mosquitoes; I_A_ infected adult mosquitoes; and W for the viral load in breeding site in PFU. The other parameters are defined in Table 1.

**Table 1.**
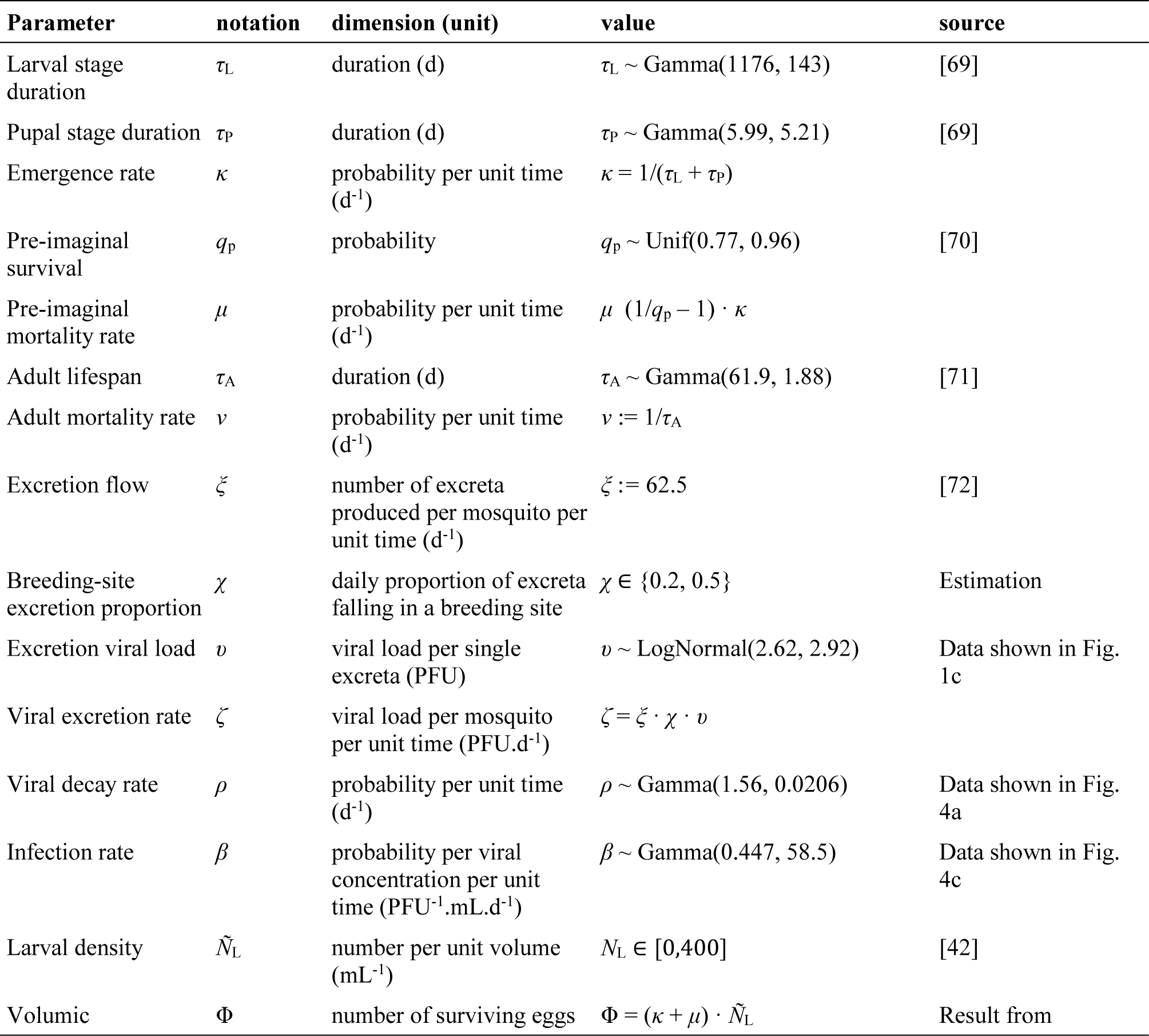
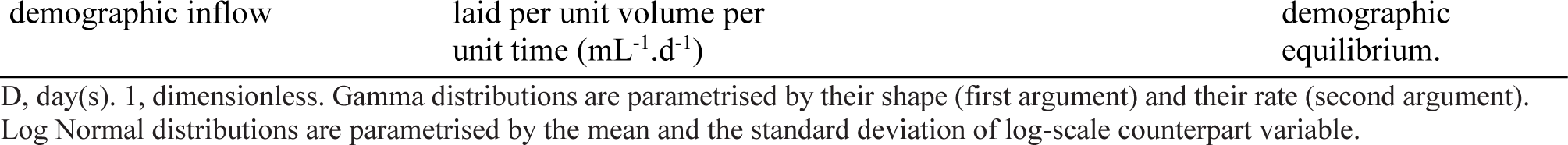
List of parameters involved in the computation of the WNV diagonal reproduction number.

The distribution of the *R*^d^ was calculated using Monte-Carlo method by computing its value across a large number (10,000) of parameter sets, independently drawn (both within and between sets) from distributions fitted from data either found in the literature or generated by the current study (Table 1). Note that the volumic demographic inflow Φ is inked to the (volumic) larval density *N*_L_, defined as the value of (*S*_L_ + *I*_L_)/*V* (i.e., the total number of larvae and pupae in the breeding site, whether susceptible or infected, per unit volume) and evaluated at the demographic equilibrium (i.e. by cancelling out the ODE 1-3).

Modelling assumptions included the well-mixed nature of the breeding site water volume, the exponential distribution of the time-to-events (conditionally to the knowledge of their expectations), the negligibility of the WNV infection impact on both immature and mature stage survival, the non-susceptibility of the eggs and the density-dependence of mosquito demography restricted by breeding site volume [in line with empirical studies suggesting fitness reduction in overcrowded habitat [32]].

All calculations and visualisations of the modelling part were performed on R [33], using the package fitdistrplus [34] for distribution fitting.

### Statistics

Differences in infection rate were tested with Chi-square. One-way repeated-measures ANOVA was used to test the effect of DPE on infection intensity. Statistical analyses were conducted with Prism v8.0 (GraphPad).

## Results

### Quantification of infectious virions in mosquito excreta

To test whether excreta from infected mosquitoes carry infectious virions, we orally infected *Cx. quinquefasciatus* with 10^5^ PFU/ml of WNV. We collected pools of excreta across different days post exposure (DPE) and inoculated virus-susceptible Vero cells with the excreta solution. At 6 days post inoculation, we detected viral genomic RNA (gRNA) in the resulting cell supernatant (Fig. 1a), demonstrating active viral infection. To confirm that excreta induced a productive infection, we performed a cell-based titration assay and showed that the supernatant of cells inoculated with mosquito excreta contained infectious virions as indicated by the presence of many lytic plaques on the cell monolayer (Fig. 1b). In contrast, cells inoculated with excreta of non-infected mosquitoes did not show any plaque. As observed in previous studies [29], our results confirm that excreta of infected mosquitoes, in our case *Cx. quinquefasciatus* mosquitoes infected with WNV, carry infectious viral particles.

**Figure 1.**
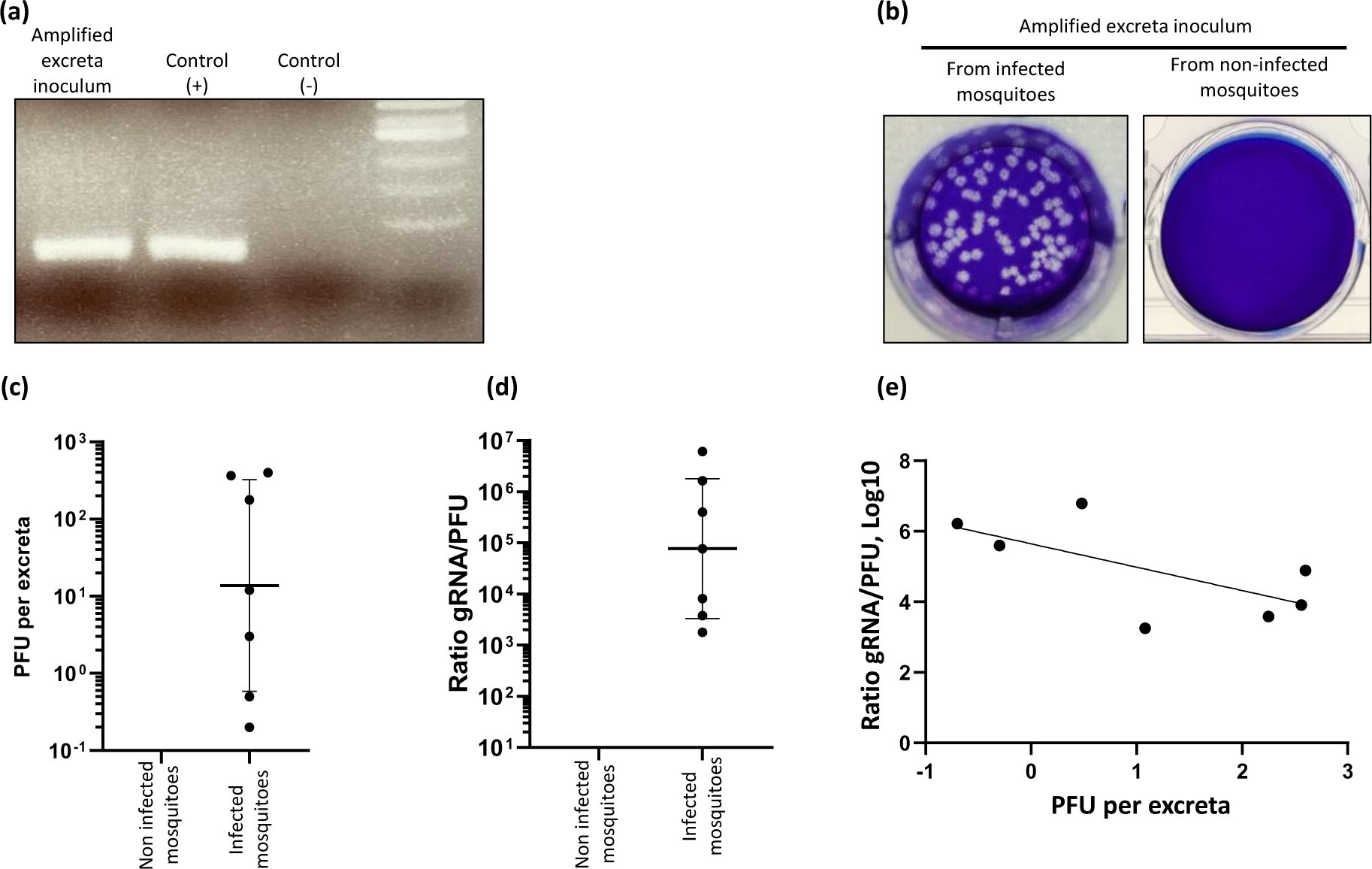
Detection and quantification of infectious viruses in mosquito excreta. **(a, b)** Detection of WNV viral RNA (a) and infectious particles (b) in supernatant from cells infected with excreta pools (i.e., amplified excreta inoculum). Control (+) corresponds to RNA extracts from WNV stock. Control (-) corresponds to water. **(c, d)** Quantification of PFU per excreta (c) and ratio of viral genomic RNA (gRNA)/PFU in the same excreta pools collected 6 days post mosquito exposure to blood containing 5 x 10^6^ PFU/ml. Bars show geometric means ± S.D. Each point indicates one excreta pool. **(e)** Correlation between PFU per excreta and gRNA/PFU ratio for the previous samples.

We next quantified the number of infectious particles per excreta. To enable excreta counting, we offered mosquitoes a sugar solution supplemented with food colorant that resulted in blue-colored excreta and counted the blue dots on the plastic walls as a proxy for excreta. To maximize the number of collected excreta, we grouped 25 mosquitoes in one container and regularly collected excreta by washing the plastic containers with cell culture media used to perform viral titration. However, we could not detect infectious particles when collection was conducted every 24h or more. We reasoned that viruses may not be stable for long time in dried excreta and collected excreta at shorter intervals of 1h-1h30 to limit virus degradation. In these conditions, we detected infectious particles in pools of excreta and were able to quantify the number of PFU, which we divided by the estimated number of excreta to obtain an averaged PFU/excreta. We observed a large variation in PFU/excreta between the different samples ranging from 0.2 to 400 PFU/excreta with a geometric mean of 13.75 PFU/excreta (Fig. 1c). As a control, we did not detect any plaque in control excreta from mosquitoes that were not exposed to an infectious blood meal.

To evaluate the infectivity of excreted virions, we calculated the ratio of gRNA/PFU, which estimates the number of infectious particles among all particles [35]. For this, we assumed that each particle contained one gRNA copy and each PFU resulted from one infectious unit. In excreta, the gRNA/PFU ratio exhibited variability, ranging from 1.8 x 10^3^ to 6.1 x 10^6^, with a geometric mean of 7.8 x 10^4^ (Fig. 1d). In comparison to a gRNA/PFU ratio of 100 for dengue virus, another flavivirus, secreted from mosquito cells [36], the higher gRNA/PFU ratio for excreted WNV indicates a high proportion of non-infectious particles, which may have undergone degradation before excreta collection. We reasoned that the elevated gRNA/PFU ratio might be attributed to virion degradation in certain samples, given the varied collection times contingent on mosquito excretion dynamics. Supporting this hypothesis, we observed a clear negative correlation (R² = 0.44) between excreta infection load, measured by PFU/excreta, and virion infectivity, estimated by gRNA/PFU ratio (Fig. 1e). This observation underscores the sensitivity of excreted virions in our conditions, implying an underestimation of PFU per excreta. Altogether, our findings demonstrate that WNV-infected mosquitoes excrete infectious virions, which quantification at an average of 13.75 PFU per excreta was probably underestimated due to virus lability.

### Virions are excreted early and continuously after exposure to an infectious blood meal

To deepen our comprehension of virion excretion, we assessed the kinetics of virion excretion and how mosquito infection level influences virion excretion. To monitor the time period of excretion, we collected excreta from single mosquitoes every 2 days from 4-12 DPE to a WNV blood inoculum of 10^7^ PFU/ml, which is within the high end of bird viremia [37,38]. Excreta collected at each time point corresponded to all excreta from the past 2 days. For instance, sample at 4 DPE included excreta from 2-4 DPE. We did not collect excreta earlier than 2 DPE to avoid collecting viruses from the blood inoculum [39]. We then quantified viral gRNA and calculated both the infection rate, as the percentage of samples with detectable amount of gRNA among collected samples, and the infection intensity, as gRNA copies per infected samples.

First, we quantified infection in the orally exposed mosquitoes from which we collected excreta at the end of the experiment (12 DPE). The high blood inoculum resulted in 100% of mosquitoes infected with a geometric mean of 3.2 x 10^8^ gRNA copies per mosquito (Fig. 2a). Second, we observed that about 50% of excreta carried viruses as early as 4 DPE and that excreta infection rate peaked at 93% at 6 DPE before gradually decreasing to 50% at 10 and 12 DPE (Fig. 2a). In contrast, the infection intensity (i.e., gRNA copies per infected samples) did not significantly change with time and remained relatively constant between 2.6 x 10^7^ and 5.5 x 10^8^ gRNA copies per excreta sample across the different time points (Fig. 2a).

**Figure 2.**
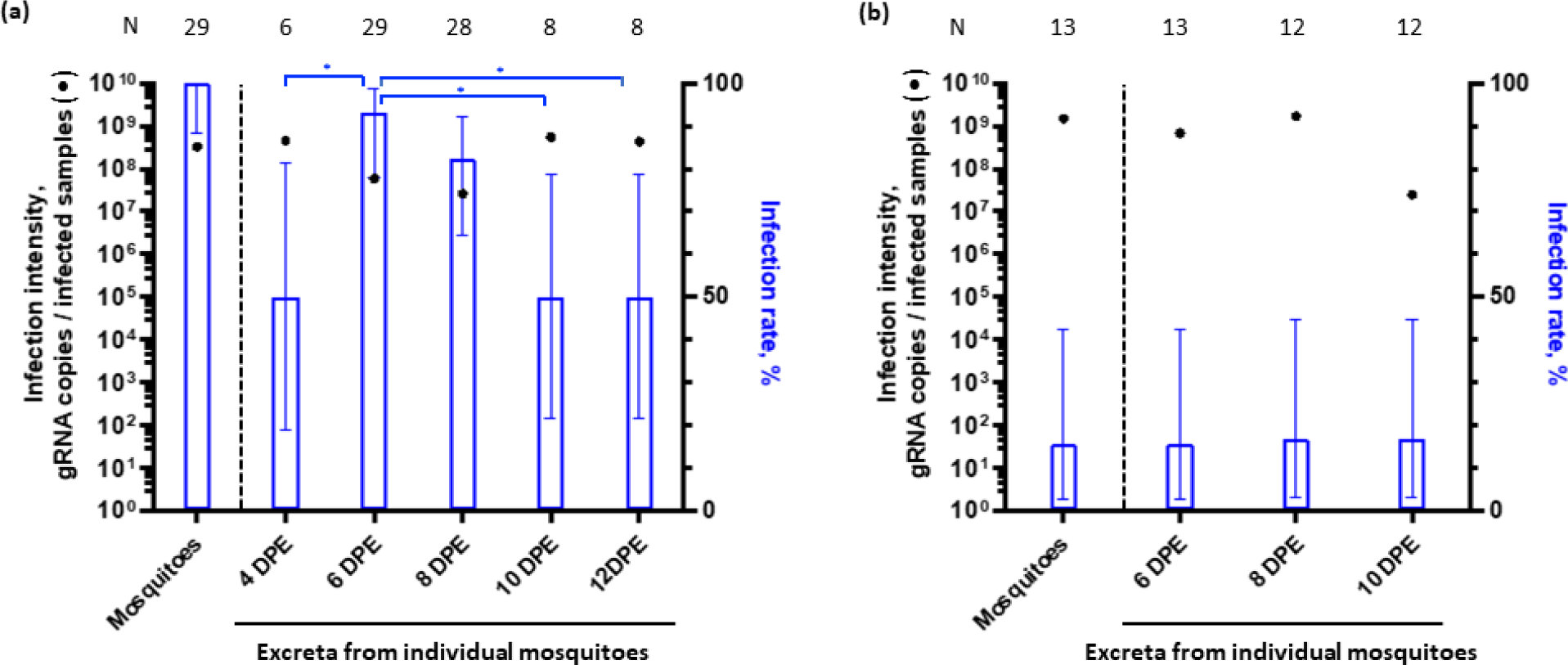
The effect of oral inoculum and days post exposure (DPE) on virus excretion. **(a, b)** Infection intensity and infection rate in mosquitoes exposed to blood containing 10^7^ (a) or 10^5^ (b) PFU/ml of WNV and in their excreta collected every two days. Black dots show geometric mean ± S.D for infection intensity. Blue bars show percentage ± 95% C.I. for infection rate. N, number of samples. Chi² was used to compare infection rates. Mixed-effects one-way ANOVA was used to compare infection intensities. *, p < 0.05.

To evaluate the influence of mosquito infection level, we repeated the excreta collection kinetics with mosquito orally exposed to a lower inoculum (i.e., 10^5^ PFU/ml) of WNV, resulting in 15% of infected mosquitoes with a geometric mean of 1.5 x 10^9^ gRNA copies per mosquito collected at 10 DPE (Fig. 2b). Excreta infection rate from 6-10 DPE was stable between 15-17% (Fig. 2b), and gRNA was mostly detected in excreta from infected-mosquitoes. Additionally, we found that each infected excreta samples contained 6.8 x 10^8^ and 1.7 x 10^9^ gRNA copies at 6 and 8 DPE, respectively, before diminishing to 2.5 x 10^7^ gRNA copies at 10 DPE (Fig. 2b). Altogether, the kinetic study from mosquitoes infected with a high and low inoculum show that virions are excreted early after oral exposure to infectious blood and for a long period at a relatively constant intensity level.

### Pupae are highly susceptible to infection

To determine whether infectious excreta can infect mosquito aquatic stages, we first monitored the virus stability in mosquito rearing water. WNV was diluted in water supplemented with larval food and quantified at different time intervals. At the initial collection time (0 min), just after diluting the virus stock, the number of infectious particles was 1,425 PFU/ml (Fig. 3a). Infectious particles then rapidly diminished to reach zero at 1h post inoculation, indicating a high lability of the virus in rearing water.

**Figure 3.**
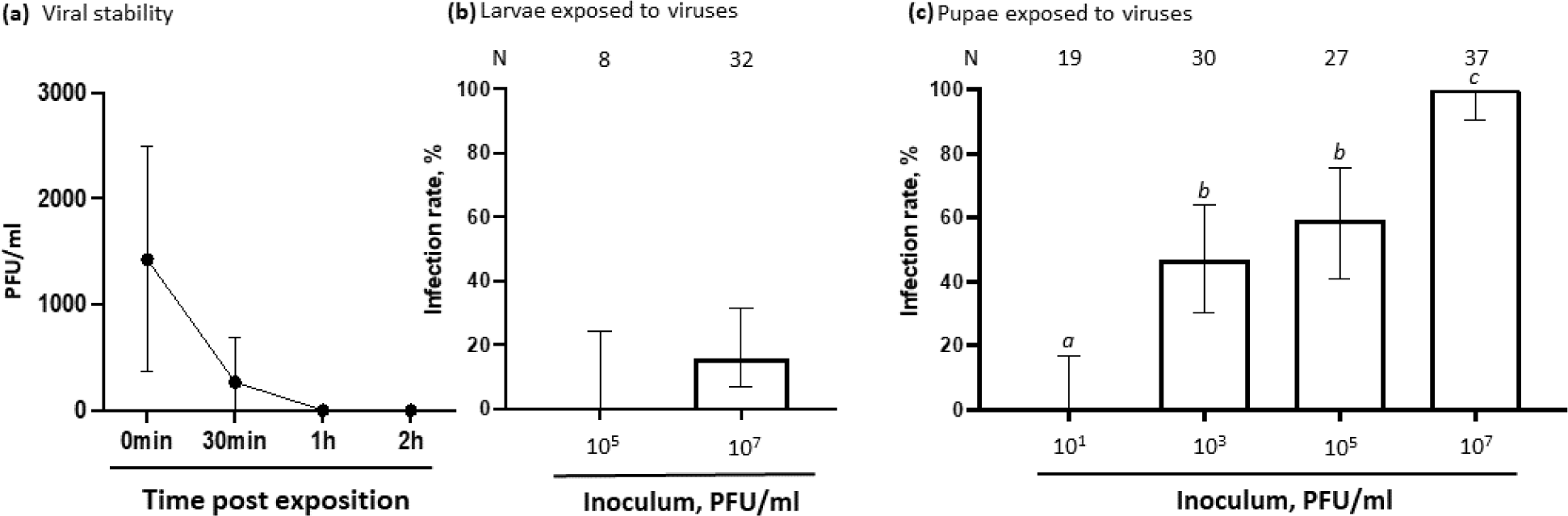
Susceptibility of aquatic stages to WNV exposure. **(a)** Stability of WNV in rearing water. Points indicate mean ± s.e.m of PFU/ml in water at different time post inoculation. N, 4. **(b, c)** Infection rate for L4 larvae exposed to WNV at L1 stage (b) and for adult mosquitoes exposed at the pupal stage (c). Bars show percentage ± 95% C.I. Chi² was used to compare infection rates between different virus concentrations. Different letters indicate significant differences, p < 0.05. N, number of individual mosquitoes.

We evaluated the susceptibility of L1 larvae and pupae to different concentrations of WNV in rearing water. Our experimental design included several precautions to avoid confounding effects. Mosquito aquatic stages were exposed for only 1h to minimize effects due to exposition to the viral stock solution. Viral gRNA was quantified in extensively washed L4 larvae resulting from the exposed L1 larvae to avoid detecting viral remnants from the inoculum. For the same reason, viral gRNA was quantified in adult mosquitoes resulting from the exposed pupae, as gut content is expelled and cuticula renewed during morphogenesis [40]. None of the larvae were infected after exposition to 10^5^ PFU/ml and only 15% after incubation with 10^7^ PFU/ml (Fig. 3b). In contrast, pupae were more susceptible to infection with 46% infected with 10^3^, 59% with 10^5^ and 100% with 10^7^ PFU/ml (Fig. 3c). We also evaluated survival after inoculum exposition. Larvae were not affected, whereas nymphs exhibited a slightly reduced survival (Sup. Fig. 2). Altogether, our results show that the short duration stability of WNV in rearing solution is sufficient to infect larvae and pupae, albeit pupae are more susceptible to infection. These observations imply that mosquito excretion in rearing water pools may lead to infection of aquatic stages.

### Infectious mosquito excreta infect pupae

While our previous experiments separately determined the excreta infectivity and the infection susceptibility of immature mosquitoes, we then tested the proof-of-concept that infectious excreta can infect mosquito pupae. We collected pools of excreta every 1h from mosquitoes at 6 DPE to a high blood inoculum to ensure maximum excreta infectivity. The excreta pools were quantified and diluted in rearing water at 4.6 x 10^3^ PFU/ml. Pupae were reared in the excreta-containing solution and infection rate assessed in adult mosquitoes. We found that 17% of pupae-exposed adults were infected (Fig. 4), thereby establishing the proof-of-concept of excreta-mediated transmission of an arbovirus.

**Figure 4.**
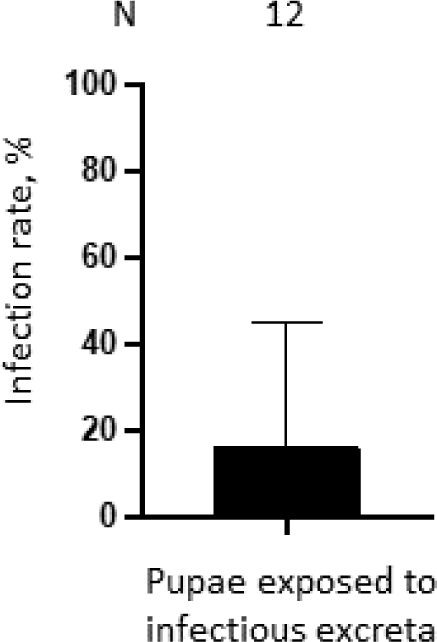
Susceptibility of pupae to infectious excreta. Bar shows infection rate + 95% C.I. in adult mosquitoes exposed at the pupal stage to infectious excreta at a concentration of 4.6 x 10^3^ PFU/ml. N, number of individual mosquitoes.

### Excreta-mediated infection can maintain flavivirus infection within mosquito populations

To determine the contribution of excreta-mediated infection in WNV epidemiology within mosquito reservoir, we built and examined a compartmental model (Fig. 5a). In a given breeding site, we modelled egg laying, mortality, hatching and emergence to calculate the number of susceptible immature mosquitoes (S_L_) and the resulting number of infected immature mosquitoes (I_L_) based on excreted virions (W) from infected adult mosquitoes (I_A_). The resulting basic reproduction number is *R*^d^_0_ = ∛(((𝑁^∼^_𝐿_) ∼𝜅𝜁𝛽)/(𝜅 + 𝜇)𝜈𝜌) (see Table 1 for details of the parameters). This formula implies that the epidemiological potential of excreta-mediated infection increases with larval density (*Ñ*_L_), survival rate to emergence (*κ/*(*κ + μ*)), excretion rate (*ζ*), infection rate (*β*), duration of the adult stage (ν^-1^) and time before excreted virions lose their infectivity (*ρ*^-1^). Importantly, the analysis of the model shows that the epidemiological potential does not depend on the surface, height or volume of the breeding sites. As a consequence, our modelling result is scale-free and applies to any size of mosquito population, or any spatial range. Moreover, our reproduction number represents solely the lower bound of the true basic reproduction number, as the model does not account for any other transmission route - namely horizontal, from mammal hosts to mosquitoes, and vertical, from female mosquitoes to eggs [41].

**Figure 5.**
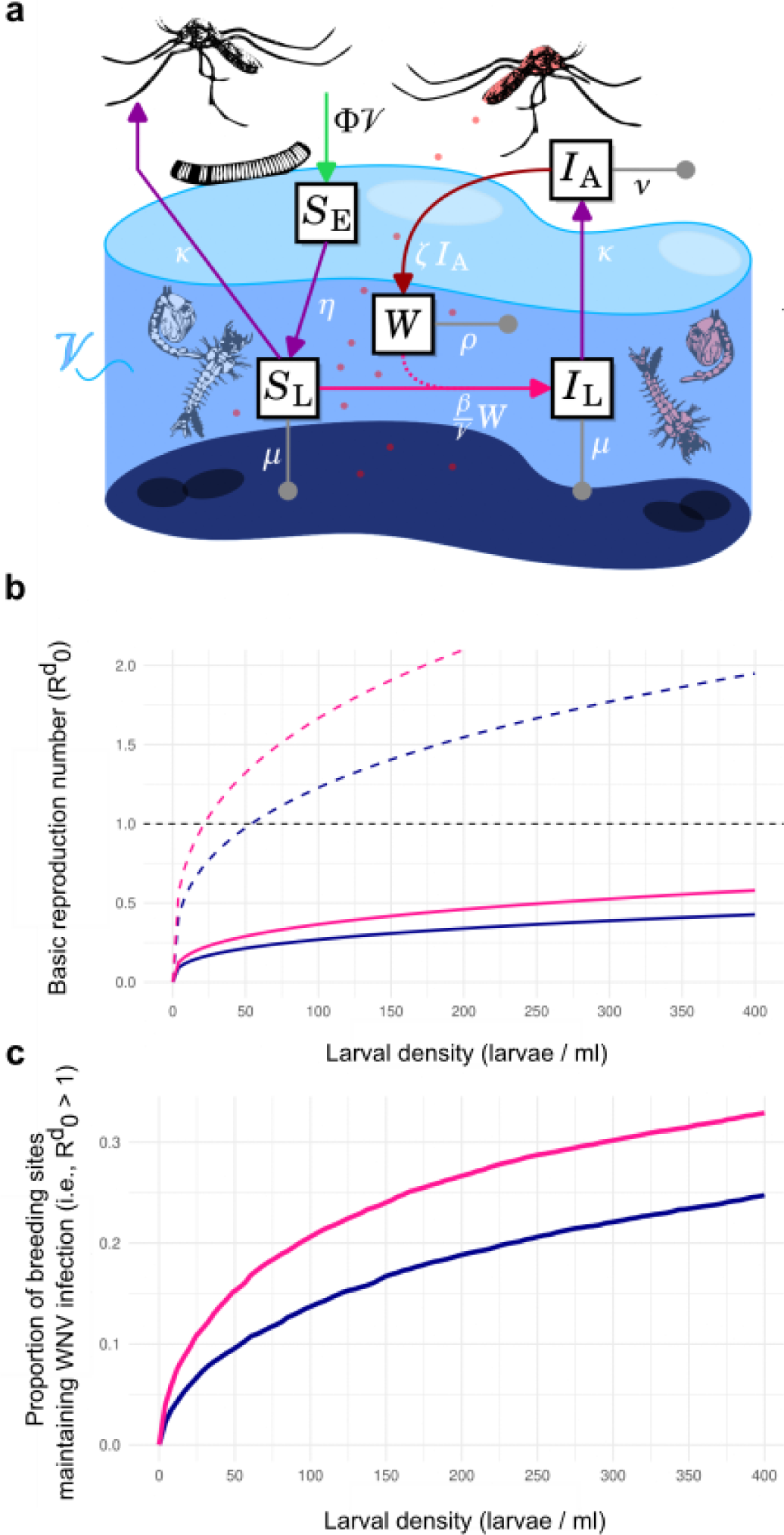
Mathematical modelling of excreta-mediated infection of mosquito aquatic stages. **(a)** Flow chart and mathematical formulation of the excreta-mediated flavivirus transmission model. V, breeding site volume. ΦV, egglaying rate. S_E_, egg survival. η, egg hatching rate. S_L_, immature mosquito susceptibility to infection. µ, immature mosquito mortality. βW/V, immature mosquito infection where W represents the WNV load in the breeding site (assumed well-mixed) and β the infection rate. I_L_, infected immature mosquitoes. κ, adult emergence. I_A_, infected adult mosquitoes. ν, adult mosquito mortality. ζ, rate of virion excretion into the breeding site. p, decay of excreted virions. Red mosquitoes indicate infection. **(b)** Basic reproduction number, *R*^d^, as a function of larval density. Solid curves indicate the median and dashed curve the 90^th^ percentile. **(c)** The proportion of breeding sites maintaining WNV infection (*R*^d^ > 1) as a function of larval density. (b-c) Blue curves indicate values for a fraction of excreta falling in a breeding site set at 20%, while pink curves indicate values for a fraction at 50%.

Feeding the model with data from our study and literature (Table 1), we inferred the distribution of *R*^d^ as a function of larval density, ranging from 0-400 larva per L – a density range previously observed in the field [42]. Although extremely hard to assess in nature, the proportion of excreta falling into breeding sites is a key determinant of *R*^d^ . In absence of data, we selected two reasonable boundaries at 20 and 50% for the proportion of excreta falling into a breeding site. The median basic reproduction number for excreta-mediated transmission rapidly increased from 0 to 25 larva/L and subsequently gradually increased to 0.42 and 0.57 for 400 larvae/L with 20 and 50% excreta falling into breeding sites, respectively (Fig. 5b). A reproduction number lower than 1 indicates that excreta-mediated transmission cannot amplify transmission. However, by computing variability in conditions between breeding sites, our model showed that the 90^th^ percentile of reproduction number reached 1 for as little as 25 and 50 larvae/L for 20 and 50% excreta falling into breeding sites, respectively. Accordingly, when plotting the proportion of breeding sites suitable for excreta-mediated infection, we calculated that transmission takes place in some breeding sites (Fig. 5c). For instance, excreta-mediated infection occurs in 14 and 20% of breeding sites containing a low density of 100 larvae/L when 20 and 50% excreta falling into breeding sites, respectively. Altogether, by combining detailed characterization of the parameters defining excreta-mediated infection of mosquitoes and comprehensive mathematical modelling, we revealed the existence of a new mode of transmission within mosquito populations through infectious excreta.

## Discussion

While mosquito-human transmission (horizontal) remains the most prevalent route, repeated detection of multiple flaviviruses, including WNV, in non-blood feeding mosquito stages such as males, larvae and pupae expose the existence of alternative modes of transmission [23–26,43–47]. In our study, we demonstrate that transmission occurs when infected mosquitoes release excreta in breeding sites. We reported the presence of infectious WNV virions in mosquito excreta and quantified a potentially high concentration of infectious units per excreta. By defining the mosquito-related conditions for virion excretion, we observed that virion excretion occurs shortly after mosquito oral infection and remain constant for longer periods. We also found that excreta viral load is independent of infection level, as previously observed [28]. These findings emphasize the infectiousness of excreta from infected mosquitoes. Furthermore, we reported the susceptibility of immature mosquitoes, especially pupae, to WNV infection, and demonstrated the capacity of infectious excreta to infect immature mosquitoes, uncovering a new mode of transmission. Finally, we modelled excreta-mediated transmission in the field and demonstrated its potential for maintaining WNV infection within mosquito reservoirs. As compared to horizontal and vertical transmissions, we propose to name the excreta-mediated transmission as “diagonal transmission”.

Excreta-mediated transmission depends on several parameters. First, infectious virions have to be shed through excretion. Malpighian tubules are the main excretory organs and accumulate wastes as primary urine, which is then transferred to the hindgut for excretion [48]. Flavivirus infection of the Malpighian tubules [28] can result in virion accumulation in urine and subsequent excretion. Alternatively, following initial infection of the midgut, virions can be secreted into the gut lumen and channeled to the hindgut for excretion. Other authors detected a very low WNV inoculum in excreta from *Cx. annulirostris*, suspecting degradation by proteases [29]. Based on our observed sensitivity to time for excreted WNV, we posit that the previously-observed low infectivity resulted from the bi-daily excreta collection. Second, mosquitoes have to drop excreta in breeding sites. Excretion in mosquitoes occurs continuously but more frequently when the insect imbibes liquids, given the osmoregulation function of excretion [48,49]. Accordingly, mosquitoes exhibit an excretion peak shortly after blood-feeding [50]. Mosquitoes drinking from breeding sites should similarly excrete in excess, contaminating the water. Additionally, excreta could be released during egg laying by compressing the hindgut. In our model, we selected a conservative and a more “optimistic” estimate of excreta proportions falling into breeding sites, both of which resulted in maintenance of WNV in certain mosquito reservoirs. Third, there must be immature mosquitoes in the breeding sites. Mosquito selection of breeding sites with specific characteristics [51,52] and attraction to breeding sites with con-specific eggs [53,54] because of egg aggregation pheromone [55,56] should favor this condition. Forth, viruses have to be stable in breeding water. WNV half-life in cell culture media is 17h [57]. We observed a much faster viral decay in laboratory breeding water that may be caused by unfavorable pH [58]. Viral stability is expected to fluctuate depending on breeding site biophysical conditions, such as pH, oxygen level, temperature and organic matter concentration. Last, immature mosquitoes have to be susceptible to infection. Although both larvae and pupae were susceptible to WNV infection, we observed a higher susceptibility for pupae. Infection of immature mosquitoes was previously observed for Zika and dengue viruses [59,60], while the differential susceptibility between larvae and pupae was also previously reported [60]. Infection may occur when viruses come in contact with midgut epithelial cells. However, larvae midgut has a protective peritrophic membrane that is absent in nymph [40,61], potentially explaining the differential susceptibility between the two immature stages. Alternatively, changes in cuticle during the nymphal stage might favor virus penetration [62]. Our results demonstrate that each of these conditions is met to allow excreta-mediated transmission.

Excreta-mediated transmission potentially occurs in all arbovirus-mosquito systems because all the required conditions are conserved in different arbovirus-mosquito systems. Multiple flaviviruses such as dengue, Usutu, Murray valley viruses [28,63,64] and alphaviruses such as Ross river virus [29] shed virions in excreta from *Aedes* and *Culex* mosquitoes, although excreta infectivity has not been tested. WNV [65] and Zika virus [60] survive in water from potential breeding sites, while all four serotypes of dengue virus remain infectious in cell media [57]. Finally, immature mosquitoes from *Aedes* and *Culex* are susceptible to Zika [60], dengue [59] and Rift valley fever viruses [66]. Importantly, conservation of excreta-mediated transmission across different arbovirus systems implies the potential for this mode of transmission to act as a transmission bridge for viruses between different mosquito vectors. Indeed, breeding sites usually contain several different mosquito species [67,68]. A shift in mosquito vectors to more anthropophilic species could promote emergence of zoonotic arboviruses.

Understanding arbovirus transmission routes is critical to deploy efficient vector control strategies. Our discovery of a new excreta-mediated (diagonal) mode of transmission emphasizes the importance of water management. Restricting excreta-mediated transmission will alleviate the arbovirus health burden by reducing maintenance of arbovirus reservoirs in mosquito populations and preventing expansion of arbovirus host range through a switch in mosquito vector species.

## Acknowledgement

MSc scholarship for QN was provided by KIM RIVE from MUSE. PhD scholarship for IS was provided by the Institut Méditerranéen Hospitalier (IHU, Marseille). Support for the research was provided by the Labex CEMEB and the I-SITE Excellence Program of the University of Montpellier, under the investissements France 2030 as the exploratory research project INGENIOUS to RH, the French Agence Nationale pour la Recherche (ANR-20-CE15-0006) and by the EU (HORIZON-HLTH-2023-DISEASE-03 #101137006) to JP.

## Supplementary Figures

**Sup. Figure 1.**
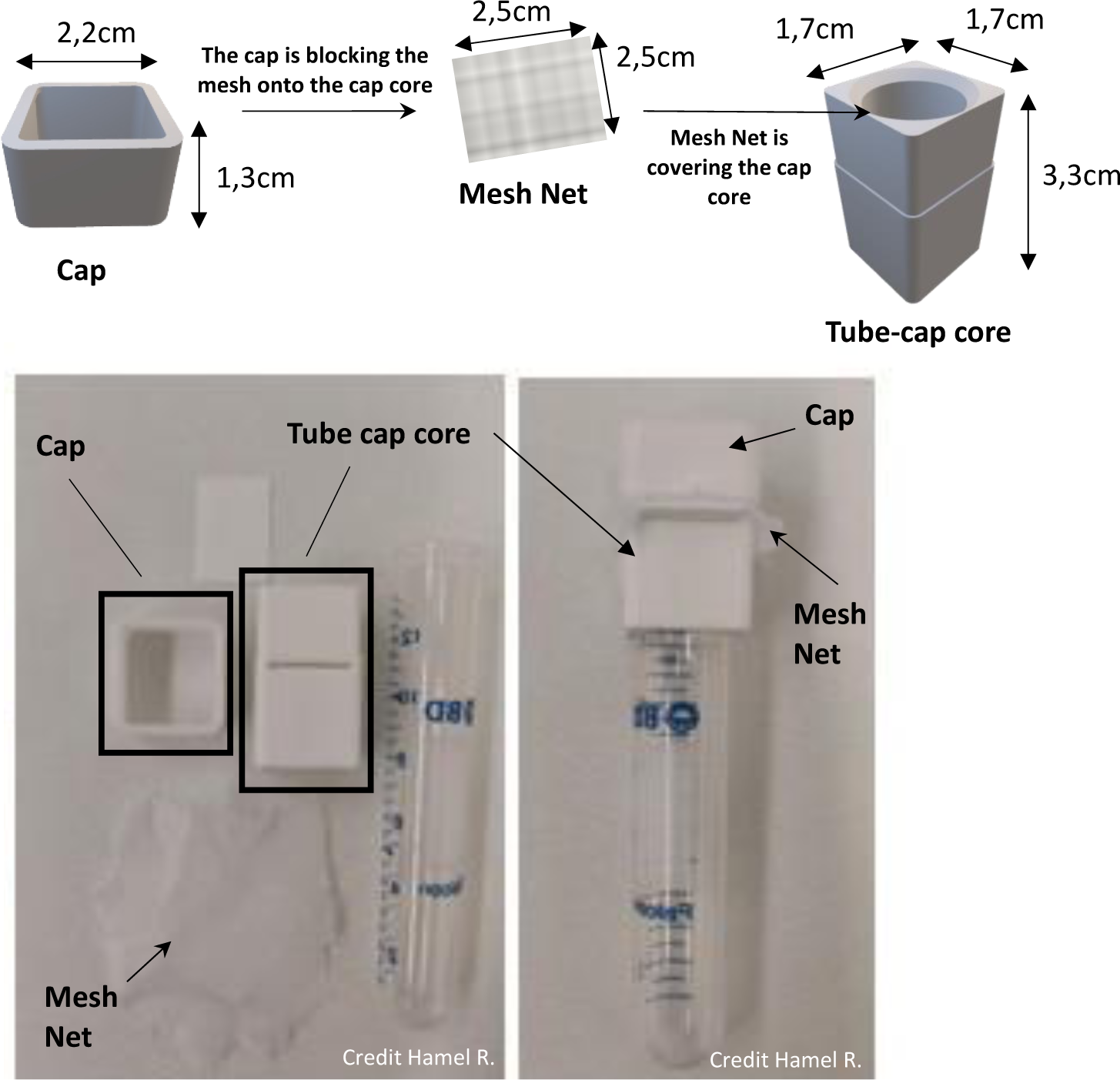
Scheme of the tube-cap manufactured to enable safe transfer of mosquitoes to other tubes. Drawing and pictures of the tube-cap. Designed by Dr. Albin Fontaine (Unité Parasitologie et Entomologie, Département Microbiologie et maladies infectieuses, Institut de Recherche Biomédicale des Armées, Marseille, France). STL files available on IRD-DataSuds.

**Sup. Figure 2.**
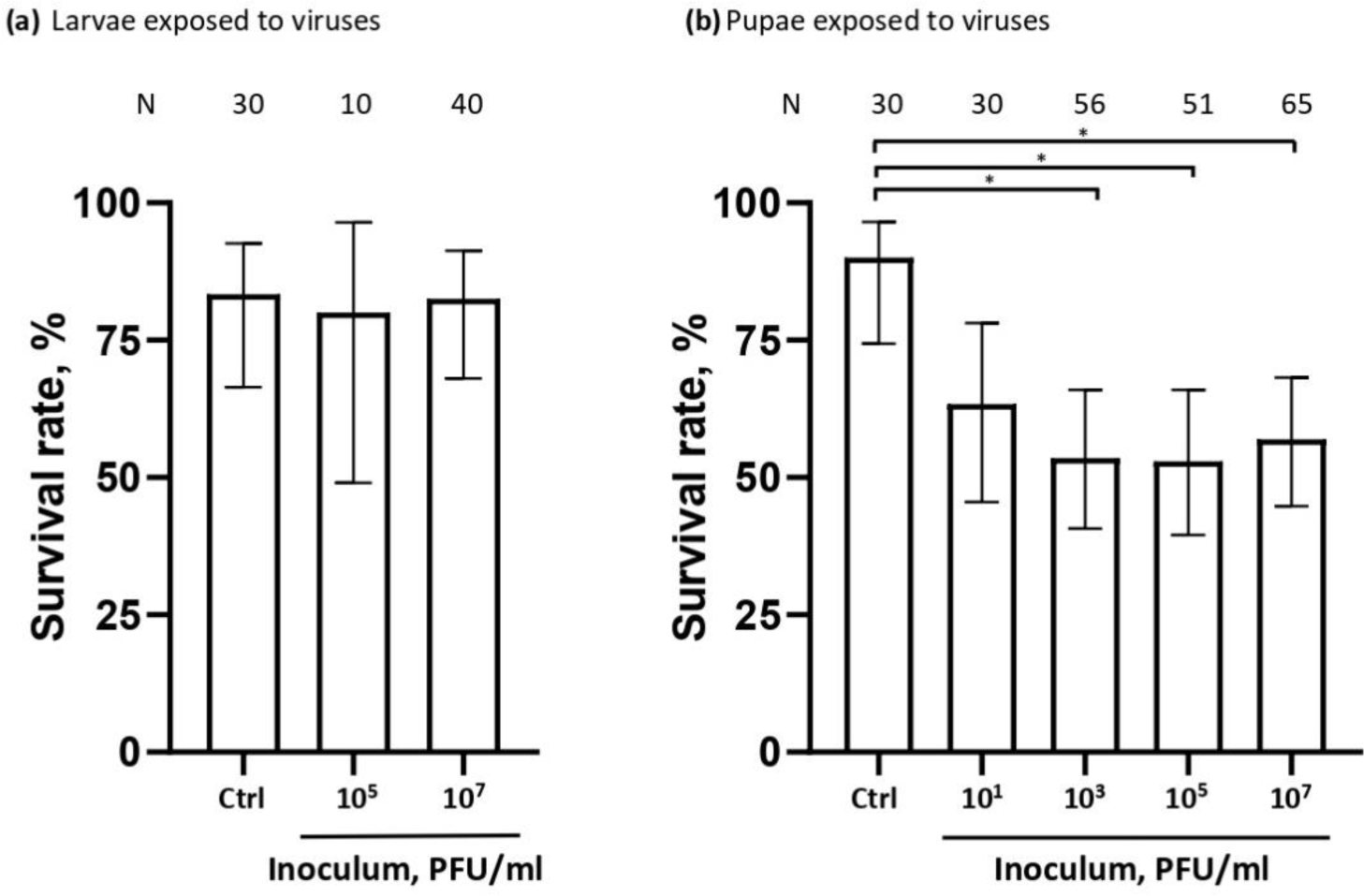
Survival of mosquito aquatic stages exposed to WNV inoculum. (**a-b**) Survival rate for L4 larvae exposed as L1 Larvae (a) and adult mosquitoes exposed as pupae (b). Bars show percent ± 95% C.I. N, number of individual mosquitoes. *, p < 0.05, as determined by χ² test.

## References

1. Smithburn KC, Hughes TP, Burke AW, Paul JH. A Neurotropic Virus Isolated from the Blood of a Native of Uganda. The American Journal of Tropical Medicine and Hygiene. 1940;s1–20: 471–492. doi:10.4269/ajtmh.1940.s1-20.471

2. Heidecke J, Schettini AL, Rocklöv J. West Nile virus eco-epidemiology and climate change. PLOS Climate. 2023;2: e0000129. doi:10.1371/journal.pclm.0000129

3. Kramer LD, Li J, Shi P-Y. West Nile virus. The Lancet Neurology. 2007;6: 171–181. doi:10.1016/S1474-4422(07)70030-3

4. Pradier S, Lecollinet S, Leblond A. West Nile virus epidemiology and factors triggering change in its distribution in Europe. 2012;31: 829. doi:10.20506/rst.31.3.2167

5. Ronca SE, Ruff JC, Murray KO. A 20-year historical review of West Nile virus since its initial emergence in North America: Has West Nile virus become a neglected tropical disease? PLOS Neglected Tropical Diseases. 2021;15: e0009190. doi:10.1371/journal.pntd.0009190

6. Zeller HG, Schuffenecker I. West Nile Virus: An Overview of Its Spread in Europe and the Mediterranean Basin in Contrast to Its Spread in the Americas. Eur J Clin Microbiol Infect Dis. 2004;23: 147–156. doi:10.1007/s10096-003-1085-1

7. Suthar MS, Diamond MS, Gale Jr M. West Nile virus infection and immunity. Nature Reviews Microbiology. 2013;11: 115–128.

8. Ciota AT, Kramer LD. Vector-Virus Interactions and Transmission Dynamics of West Nile Virus. Viruses. 2013;5: 3021–3047. doi:10.3390/v5123021

9. Historic Data (1999-2022) | West Nile Virus | CDC. 13 Jun 2023 [cited 4 Sep 2023]. Available: https://www.cdc.gov/westnile/statsmaps/historic-data.html

10. Mandalakas AM, Kippes C, Sedransk J, Kile JR, Garg A, McLeod J, et al. West Nile Virus Epidemic, Northeast Ohio, 2002. Emerging Infectious Diseases. 2005;11: 1774–7. doi:10.3201/eid1111.040933

11. Epidemiological update: West Nile virus transmission season in Europe, 2022. 30 Jun 2023 [cited 1 Sep 2023]. Available: https://www.ecdc.europa.eu/en/news-events/epidemiological-update-west-nile-virus-transmission-season-europe-2022

12. Chancey C, Grinev A, Volkova E, Rios M. The Global Ecology and Epidemiology of West Nile Virus. In: BioMed Research International [Internet]. Hindawi; 19 Mar 2015 [cited 6 Mar 2021] p. e376230. 10.1155/2015/376230

13. Farajollahi A, Fonseca DM, Kramer LD, Marm Kilpatrick A. “Bird biting” mosquitoes and human disease: a review of the role of Culex pipiens complex mosquitoes in epidemiology. Infection Genetics and Evolution. 2011;11: 1577–85.

14. Molaei G, Andreadis TG, Armstrong PM, Bueno R, Dennett JA, Real SV, et al. Host feeding pattern of Culex quinquefasciatus (Diptera: Culicidae) and its role in transmission of West Nile virus in Harris County, Texas. The American journal of tropical medicine and hygiene. 2007;77: 73–81.

15. Komar N, Langevin S, Hinten S, Nemeth N, Edwards E, Hettler D, et al. Experimental infection of North American birds with the New York 1999 strain of West Nile virus. Emerg Infect Dis. 2003;9: 311–322. doi:10.3201/eid0903.020628

16. McLean RG, Ubico SR, Docherty DE, Hansen WR, Sileo L, McNamara TS. West Nile virus transmission and ecology in birds. Ann N Y Acad Sci. 2001;951: 54–57. doi:10.1111/j.1749-6632.2001.tb02684.x

17. van der Meulen KM, Pensaert MB, Nauwynck HJ. West Nile virus in the vertebrate world. Arch Virol. 2005;150: 637–657. doi:10.1007/s00705-004-0463-z

18. Anderson JF, Main AJ, Delroux K, Fikrig E. Extrinsic Incubation Periods for Horizontal and Vertical Transmission of West Nile Virus by Culex pipiens pipiens (Diptera: Culicidae). Journal of medical entomology. 2008.

19. Baqar S, Hayes CG, Murphy JR, Watts DM. Vertical transmission of West Nile virus by Culex and Aedes species mosquitoes. Am J Trop Med Hyg. 1993;48: 757–762. doi:10.4269/ajtmh.1993.48.757

20. Dohm DJ, Sardelis MR, Turell MJ. Experimental vertical transmission of West Nile virus by Culex pipiens (Diptera: Culicidae). J Med Entomol. 2002;39: 640–644. doi:10.1603/0022-2585-39.4.640

21. Nelms BM, Fechter-Leggett E, Carroll BD, Macedo P, Kluh S, Reisen WK. Experimental and natural vertical transmission of West Nile virus by California Culex (Diptera: Culicidae) mosquitoes. J Med Entomol. 2013;50: 371–378. doi:10.1603/me12264

22. Manuel M, Missé D, Pompon J. Highly Efficient Vertical Transmission for Zika Virus in Aedes aegypti after Long Extrinsic Incubation Time. Pathogens. 2020;9: E366. doi:10.3390/pathogens9050366

23. Miller BR, Nasci RS, Godsey MS, Savage HM, Lutwama JJ, Lanciotti RS, et al. First field evidence for natural vertical transmission of West Nile virus in Culex univittatus complex mosquitoes from Rift Valley province, Kenya. Am J Trop Med Hyg. 2000;62: 240–246. doi:10.4269/ajtmh.2000.62.240

24. Phillips RA, Christensen K. Field-Caught Culex erythrothorax Larvae Found Naturally Infected with West Nile Virus in Grand County, Utah. moco. 2006;22: 561–562. doi:10.2987/8756-971X(2006)22[561:FCELFN]2.0.CO;2

25. Unlu I, Mackay AJ, Roy A, Yates MM, Foil LD. Evidence of vertical transmission of West Nile virus in field-collected mosquitoes. Journal of Vector Ecology. 2010;35: 95–99. doi:10.1111/j.1948-7134.2010.00064.x

26. Kolodziejek J, Seidel B, Jungbauer C, Dimmel K, Kolodziejek M, Rudolf I, et al. West Nile Virus Positive Blood Donation and Subsequent Entomological Investigation, Austria, 2014. PLOS ONE. 2015;10: e0126381. doi:10.1371/journal.pone.0126381

27. Reisen WK, Fang Y, Lothrop HD, Martinez VM, Wilson J, O’Connor P, et al. Overwintering of West Nile Virus in Southern California. Journal of Medical Entomology. 2006;43: 344–355. doi:10.1093/jmedent/43.2.344

28. Fontaine A, Jiolle D, Moltini-Conclois I, Lequime S, Lambrechts L. Excretion of dengue virus RNA by Aedes aegypti allows non-destructive monitoring of viral dissemination in individual mosquitoes. Sci Rep. 2016;6: 24885. doi:10.1038/srep24885

29. Ramírez AL, Hall-Mendelin S, Doggett SL, Hewitson GR, McMahon JL, Ritchie SA, et al. Mosquito excreta: A sample type with many potential applications for the investigation of Ross River virus and West Nile virus ecology. PLOS Neglected Tropical Diseases. 2018;12: e0006771. doi:10.1371/journal.pntd.0006771

30. Bahuon C, Desprès P, Pardigon N, Panthier J-J, Cordonnier N, Lowenski S, et al. IS-98-ST1 West Nile Virus Derived from an Infectious cDNA Clone Retains Neuroinvasiveness and Neurovirulence Properties of the Original Virus. PLOS ONE. 2012;7: e47666. doi:10.1371/journal.pone.0047666

31. van den Driessche P, Watmough J. Reproduction numbers and sub-threshold endemic equilibria for compartmental models of disease transmission. Mathematical Biosciences. 2002;180: 29–48. doi:10.1016/S0025-5564(02)00108-6

32. Ukubuiwe AC, Ojianwuna CC, Olayemi IK, Arimoro FO, Omalu ICJ, Ukubuiwe CC, et al. Quantifying the Influence of Larval Density on Disease Transmission Indices in Culex quinquefasciatus, the Major African Vector of Filariasis. Int J Insect Sci. 2019;11: 1179543319856022. doi:10.1177/1179543319856022

33. R Core Team. R: A language and environment for statistical computing. R Foundation for Statistical Computing, Vienna, Austria.; 2023. Available: https://www.R-project.org/.

34. Delignette-Muller ML, Dutang C. fitdistrplus: An R Package for Fitting Distributions. Journal of Statistical Software. 2015;64: 1–34. doi:10.18637/jss.v064.i04

35. Sanjuán R. Collective properties of viral infectivity. Current Opinion in Virology. 2018;33: 1–6. doi:10.1016/j.coviro.2018.06.001

36. Vial T, Tan W-L, Deharo E, Missé D, Marti G, Pompon J. Mosquito metabolomics reveal that dengue virus replication requires phospholipid reconfiguration via the remodeling cycle. Proc Natl Acad Sci USA. 2020; 202015095. doi:10.1073/pnas.2015095117

37. Bingham J, Lunt R, Green D, Davies K, Stevens V, Wong F. Experimental studies of the role of the little raven (Corvus mellori) in surveillance for West Nile virus in Australia. Australian Veterinary Journal. 2010;88: 204–210. doi:10.1111/j.1751-0813.2010.00582.x

38. Nemeth N, Young G, Ndaluka C, Bielefeldt-Ohmann H, Komar N, Bowen R. Persistent West Nile virus infection in the house sparrow (Passer domesticus). Arch Virol. 2009;154: 783–789. doi:10.1007/s00705-009-0369-x

39. Gooding RH, Cheung A, Rolseth B. Digestive processes of haematophagous insects. I. A literature review. 1973.

40. Moncayo AC, Lerdthusnee K, Leon R, Robich RM, Romoser WS. Meconial Peritrophic Matrix Structure, Formation, and Meconial Degeneration in Mosquito Pupae/Pharate Adults: Histological and Ultrastructural Aspects. Journal of Medical Entomology. 2005;42: 939–944. doi:10.1093/jmedent/42.6.939

41. Sofonea MT, Aldakak L, Boullosa LFV. v., Alizon S. Can Ebola virus evolve to be less virulent in humans? Journal of Evolutionary Biology. 2018;31: 382–392. doi:10.1111/jeb.13229

42. Amara Korba R, Alayat MS, Bouiba L, Boudrissa A, Bouslama Z, Boukraa S, et al. Ecological differentiation of members of the Culex pipiens complex, potential vectors of West Nile virus and Rift Valley fever virus in Algeria. Parasites & Vectors. 2016;9: 455. doi:10.1186/s13071-016-1725-9

43. Ahmad R, Ismail A, Saat Z, Lim LH. Detection of dengue virus from field Aedes aegypti and Aedes albopictus adults and larvae. Southeast Asian J Trop Med Public Health. 1997;28: 138–142.

44. Cecílio SG, Júnior WFS, Tótola AH, de Brito Magalhães CL, Ferreira JMS, de Magalhães JC. Dengue virus detection in Aedes aegypti larvae from southeastern Brazil. J Vector Ecol. 2015;40: 71–74. doi:10.1111/jvec.12134

45. Izquierdo-Suzán M, Zárate S, Torres-Flores J, Correa-Morales F, González-Acosta C, Sevilla-Reyes EE, et al. Natural Vertical Transmission of Zika Virus in Larval Aedes aegypti Populations, Morelos, Mexico. Emerg Infect Dis. 2019;25: 1477–1484. doi:10.3201/eid2508.181533

46. Johari NA, Toh SY, Voon K, Lim PKC. Detection of Zika virus RNA in Aedes aegypti and Aedes albopictus larvae in Klang Valley, Peninsular Malaysia. Trop Biomed. 2019;36: 310–314.

47. Velandia-Romero ML, Olano VA, Coronel-Ruiz C, Cabezas L, Calderon-Pelaez MA, Castellanos JE, et al. Dengue virus detection in Aedes aegypti larvae and pupae collected in rural areas of Anapoima, Cundinamarca, Colombia. Biomedica. 2017;37: 193–200.

48. Farina P, Bedini S, Conti B. Multiple Functions of Malpighian Tubules in Insects: A Review. Insects. 2022;13: 1001. doi:10.3390/insects13111001

49. Clements AN. Biology Of Mosquitoes : Development Nutrition And Reproduction By A. N. Clements. 1992.

50. Williams JC, Hagedorn HH, Beyenbach KW. Dynamic changes in flow rate and composition of urine during the post-bloodmeal diuresis inAedes aegypti (L.). J Comp Physiol B. 1983;153: 257–265. doi:10.1007/BF00689629

51. Beehler JW, Millar JG, Mulla MS. Field evaluation of synthetic compounds mediating oviposition inCulex mosquitoes (Diptera: Culicidae). J Chem Ecol. 1994;20: 281–291. doi:10.1007/BF02064436

52. Ikeshoji T, Umino T, Hirakoso S. Studies on mosquito attractants and stimulants. Part IV. An agent producing stimulative effects for oviposition of Culex pipiens fatigans in field water and the stimulative effects of various chemicals. Jikken Igaku Zasshi = Japanese Journal of Experimental Medicine. 1967;37: 61–9.

53. Bruno DW, Laurence BR. The Influence of the Apical Droplet of Culex Egg Rafts on Oviposition of Culex Pipiens Faticans (Diptera: Culicidae). Journal of Medical Entomology. 1979;16: 300–305. doi:10.1093/jmedent/16.4.300

54. Reiskind MH, Wilson ML. Culex restuans (Diptera: Culicidae) Oviposition Behavior Determined by Larval Habitat Quality and Quantity in Southeastern Michigan. Journal of Medical Entomology. 2004;41: 179–186. doi:10.1603/0022-2585-41.2.179

55. Mboera LEG, Mdira KY, Salum FM, Takken W, Pickett JA. Influence of Synthetic Oviposition Pheromone and Volatiles from Soakage Pits and Grass Infusions Upon Oviposition Site-Selection of Culex Mosquitoes in Tanzania. J Chem Ecol. 1999;25: 1855–1865. doi:10.1023/A:1020933800364

56. Millar JG, Chaney JD, Beehler JW, Mulla MS. Interaction of the Culex quinquefasciatus egg raft pheromone with a natural chemical associated with oviposition sites. J Am Mosq Control Assoc. 1994;10: 374–379.

57. Goo L, Dowd KA, Smith ARY, Pelc RS, DeMaso CR, Pierson TC. Zika Virus Is Not Uniquely Stable at Physiological Temperatures Compared to Other Flaviviruses. mBio. 2016;7: 10.1128/mbio.01396-16. doi:10.1128/mbio.01396-16

58. Oliveira ERA, de Alencastro RB, Horta BAC. New insights into flavivirus biology: the influence of pH over interactions between prM and E proteins. J Comput Aided Mol Des. 2017;31: 1009–1019. doi:10.1007/s10822-017-0076-8

59. Bara JJ, Clark TM, Remold SK. Susceptibility of Larval Aedes aegypti and Aedes albopictus (Diptera: Culicidae) to Dengue Virus. Journal of Medical Entomology. 2013;50: 179–184. doi:10.1603/ME12140

60. Du S, Liu Y, Liu J, Zhao J, Champagne C, Tong L, et al. Aedes mosquitoes acquire and transmit Zika virus by breeding in contaminated aquatic environments. Nature Communications. 2019;10: 1324. doi:10.1038/s41467-019-09256-0

61. Edwards MJ, Jacobs-Lorena M. Permeability and disruption of the peritrophic matrix and caecal membrane from Aedes aegypti and Anopheles gambiae mosquito larvae. Journal of Insect Physiology. 2000;46: 1313–1320. doi:10.1016/S0022-1910(00)00053-6

62. Romoser WS, Oviedo MVN. Function of the dendritic setae in Aedes aegypti mosquito pupae: float hairs don&#39;t float. OAIP. 2011;3: 7–11. doi:10.2147/OAIP.S13727

63. Körsten C, Vasić A, Al-Hosary AA, Tews BA, Răileanu C, Silaghi C, et al. Excretion Dynamics of Arboviruses in Mosquitoes and the Potential Use in Vector Competence Studies and Arbovirus Surveillance. Trop Med Infect Dis. 2023;8: 410. doi:10.3390/tropicalmed8080410

64. Meyer DB, Ramirez AL, van den Hurk AF, Kurucz N, Ritchie SA. Development and Field Evaluation of a System to Collect Mosquito Excreta for the Detection of Arboviruses. Journal of Medical Entomology. 2019;56: 1116–1121. doi:10.1093/jme/tjz031

65. Lund M, Shearn-Bochsler V, Dusek RJ, Shivers J, Hofmeister E. Potential for Waterborne and Invertebrate Transmission of West Nile Virus in the Great Salt Lake, Utah. Applied and Environmental Microbiology. 2017;83: e00705–17. doi:10.1128/AEM.00705-17

66. Turell MJ, Linthicum KJ, Beaman JR. Transmission of Rift Valley Fever Virus by Adult Mosquitoes after Ingestion of Virus as Larvae. The American Journal of Tropical Medicine and Hygiene. 1990;43: 677–680. doi:10.4269/ajtmh.1990.43.677

67. Carrieri M, Bacchi M, Bellini R, Maini S. On the Competition Occurring Between Aedes albopictus and Culex pipiens (Diptera: Culicidae) in Italy. Environmental Entomology. 2003;32: 1313–1321. doi:10.1603/0046-225X-32.6.1313

68. Okiwelu S, Noutcha AE. Breeding Sites of Culex quinquefasciatus (Say) during the Rainy Season in Rural Lowland Rainforest, Rivers State, Nigeria. Public Health Research. 2012;2: 64–68. doi:10.5923/j.phr.20120204.01

69. Eastwood G, Cunningham AA, Kramer LD, Goodman SJ. The vector ecology of introduced Culex quinquefasciatus populations, and implications for future risk of West Nile virus emergence in the Galápagos archipelago. Med Vet Entomol. 2019;33: 44–55. doi:10.1111/mve.12329

70. Ciota AT, Matacchiero AC, Kilpatrick AM, Kramer LD. The Effect of Temperature on Life History Traits of Culex Mosquitoes. Journal of Medical Entomology. 2014;51: 55–62. doi:10.1603/ME13003

71. Moser SK, Barnard M, Frantz RM, Spencer JA, Rodarte KA, Crooker IK, et al. Scoping review of Culex mosquito life history trait heterogeneity in response to temperature. Parasites & Vectors. 2023;16: 200. doi:10.1186/s13071-023-05792-3

72. Pilotte N, Zaky WI, Abrams BP, Chadee DD, Williams SA. A Novel Xenomonitoring Technique Using Mosquito Excreta/Feces for the Detection of Filarial Parasites and Malaria. PLoS Negl Trop Dis. 2016;10: e0004641. doi:10.1371/journal.pntd.0004641

